# Comparative analyses and phylogenetic relationships of 15 *Trapa* (Trapaceae) species based on Complete Chloroplast Genomes

**DOI:** 10.1101/2021.05.17.444415

**Authors:** Xiangrong Fan, Wuchao Wang, Godfrey K. Wagutu, Wei Li, Xiuling Li, Yuanyuan Chen

**Author notes:** Correspondence: Yuanyuan Chen, Key Laboratory of Aquatic Botany and Watershed Ecology, Wuhan Botanical Garden, Center of Plant Ecology, Core Botanical Gardens, Chinese Academy of Sciences, Wuhan 430074, P. R. China., Xiuling Li, College of Life Science, Linyi University, Linyi 276000, Shandong, P. R. China. The authors are equally contributing to the study (Xiangrong Fan, Wuchao Wang).

## Abstract

*Trapa* L. is floating-leaved aquatic plants with important economic and ecological values. However, species identification and phylogenetic relationship are still unresolved for *Trapa*. In this study, complete chloroplast genomes of 13 *Trapa* species/taxa were sequenced and annotated. Combined with released sequences of the other two species, comparative analysis of cp genomes was first performed on the 15 *Trapa* species/taxa. The 15 cp genomes exhibited quadripartite structures with medium size of 155, 453-155, 559 bp. IR/SC junctions were conservative with no obvious change found. Long repetitive repeats and SSRs were mostly detected in the intergenic and LSC regions, providing useful plastid markers for species and relationship identification. Three phylogenetic analyses (MP, ML and BI) consistently showed two clusters within *Trapa*, including large- and small-seed species/taxa, respectively. This study provided the baseline information for phylogeography of *Trapa*, which would facilitate the management and utilization of genetic resources of the genus.

## 1. Introduction

Trapaceae, containing the only genus *Trapa*, is annual floating-leaved aquatic herbs naturally distributed in tropical, subtropical and temperate regions of Eurasia and Africa, and invading North America and Australia (Chen et al., 2007). APG II (The Angiosperm Phylogeny Group, 2003) equated Trapaceae with Lythraceae. However, a handful of morphological differences exist between the two families. For example, flowers of Trapaceae are solitary, 4-merous and actinomorphic, with half-inferior and slightly perigynous ovaries; Lythraceae has racemes or cymes, and the flowers are usually 4-, 6- or 8-merous, regular or irregular, with obvious perigynous ovaries. Therefore, Trapaceae is still be used today by some researchers (Chen et al., 2007). *Trapa* has important edible value because high content of starch of seeds, which has been widely cultivated as an important aquatic crop in China and India (Hummel and Kiviat, 2004). *Trapa* seed pericarps were traditional herb medicines in China, and recent studies found that the extraction of seed pericarps had bioactive components to restrain cancer, atherosclerosis, inflammation and oxidation (Akao et al., 2013; Ciou et al., 2008; Yu and Shen, 2015; Li et al., 2017; Kauser et al., 2018). Additionally, *Trapa* plants can be used to purify water body due to their excellent performance in absorbing heavy metals and nutrients (Sweta et al., 2015; Xu et al., 2020). Therefore, *Trapa* has important economic and ecological values. However, many *Trapa* species are becoming endangered or even locally extinct due to human interferences in Europe and China (Gupta and Beentje, 2017; Chen et al., field observations). Conversely, *Trapa* plants were notorious intruders in Canada and the north-eastern United States.

The knowledge of accurate species identification and phylogenetic relationship among species is essential to effectively protect and utilize genetic resources of wild plants and to manage and remove invasive plants (Campbell et al., 2009; Chorak et al., 2019). However, due to the large range of morphological variation and the lack of effective diagnostic criteria, researchers have held sharp different opinions on taxonomic classification of *Trapa*, with 1 or 2 polymorphic species or more than 20, 30 or 70 species within the genus (Cook, 1996; Vassiljev, 1949, 1965; Tutin, 1968; Yan, 1983; Kak, 1988; Chen et al., 2007). Meanwhile, the phylogenetic relationship for *Trapa* species was unresolved by the efforts of pollen morphology, cytology, quantitative classification and ontogenesis (Diao et al., 1990; Ding et al., 1991; Huang et al., 1996; Wang and Ding, 1997; Fan et al., 2016). For example, Xiong et al. (1985a, b) proposed that *T. acornis* Nakano and *T. bispinosa* Roxb were closely related based on the results of quantitative classification, while Ding et al. (1991) showed that instead of *T. bispinosa*, *T. quadrispinosa* Roxb was closed related to *T. acornis* based on their pollen morphology. And studies of quantitative classification showed that the *Trapa* species/taxa with seeds of similar size evolved closely (Xiong et al., 1990; Fan et al., 2016), which was proved by the results of allozymes (Takano and Kadono, 2005). Additionally, molecular methods further illustrated that the *Trapa* species with small seeds, *T. incisa* and *T. maximowiczii*, were of basal classification status within the genus (Jiang and Ding, 2004; Fan et al., 2021). For the large-seed *Trapa* species, a close relationship was found between *T. bispinosa* and *T. quadrispinosa* based on RAPDs and AFLPs (Jiang and Ding, 2004; Li et al., 2017; Fan et al., 2021). It is worth noting that there are two diversity centers in *Trapa* genus, one distributed in the mid-lower reaches of Yangtze River, and the other one located in the Tumen River and Amur River basins from bordering region of Russia and China (Xue et al., 2016, 2017). However, most previous studies involved fewer species/taxa, mostly collected from the mid-lower reaches of Yangtze River, and genotyped by molecular markers or nuclear genome sequencing (Li et al., 2017; Fan et al., 2021). Researches about chloroplast genome were rarely involved (Xue et al., 2017; Sun et al., 2020).

Chloroplast (cp) is a core organelle in plants for photosynthesis (Howe et al., 2003). In angiosperms, the cp genome is haploid, maternal-inherited and shaped into a DNA circle with conserved quadripartite structure. Additionally, the cp genome has slow evolutionary rate, high copy numbers per cell and compact size with 120-170kb in length (Ruhlman and Jansen, 2014). Those characteristics of cp genome, combined with the development of high-throughput technology, make sequencing of the complete cp genome an ideal tool for species identification and plant phylogenic evaluation (Cauz-Santos et al., 2017; Hong et al., 2020; Yang et al., 2018). To date, whole sequences of six *Trapa* chloroplast genomes (*T. natans*, *T. bicornis*, *T. quadrispinosa*, *T. kozhevnikovirum*, *T. incisa* and *T. maximowiczii*) have been published, but there have been no researches involving comparative analysis of the *Trapa* cp genomes. For *Trapa*, more efforts should be made to analyze the interspecific difference and assess the phylogenic relation based on the complete cp sequences.

Besides the wild species, *Trapa* also has a large number of cultivated species or varieties because of its long history of cultivation. The comprehensive analysis of cp genomes focused on wild *Trapa*, and the cultivated species or varieties would be included in another research. In this study, we sequenced the whole cp genomes of 13 wild *Trapa* species/taxa. Combined with the two published cp genomes, comprehensive analysis was first carried out within *Trapa* genus based on the 15 cp genomes data. Our specific goals were as follows: (1) to compare the chloroplast structures within *Trapa*; (2) to detect the highly variable regions as DNA barcoding markers for *Trapa* species identification; (3) to infer the phylogenetic relationship among *Trapa* species. This study will provide the baseline information for phylogeography within *Trapa* and facilitate the management and utilization of genetic resources of *Trapa*.

## 2. Materials and methods

### 2.1 Plant Materials and DNA Extraction

In the autumns of 2018 and 2019, young and healthy leaves were collected from 13 *Trapa* species/taxa. Among them, 10 species/taxa were recorded in Chinese Flora Republicae Popularis Sinicae (*T. bispinosa, T. quadrispinosa, T. japonica, T. mammillifera, T. macropoda* var. *bispinosa, T. litwinowii, T. arcuata, T. pseudoincisa, T. manshurica*, and *T. maximowiczii*; Wan, 2000); two species (*T. potaninii and T. sibirica*) were first recorded in Floral of USSR (Vassiljev, 1949). A new *Trapa* taxon was collected from Baidang Lake, Anhui province, China. The seeds of the taxon have two horns with the height from 23.4 to 34.8 mm and the width from 13.4 to 18.3 mm. The horns of the new taxon were wide and drooping, looking like a pig’s ears. Based on the nut size and the sample site, the taxon was named *Trapa natans* var. *baidangensis.* All voucher specimens were deposited in the herbarium of Wuhan Botanical Garden (HIB; Table 1). The fresh leaves were cleaned and dried in silica gel immediately. Genomic DNA was extracted from the dried leaves according to the CTAB protocol described by Doyle (1987). The DNA concentration and quality was quantified by the NanoDrop 2000 micro spectrophotometer (Thermo Fisher Scientific).

### 2.2 Chloroplast Genome Sequencing and Assembling

High quality DNA was used to build the genomic libraries. Sequencing was performed using paired end 150bp (average short-insert about 350 bp) on Illumina 2500 (Illumina, San Diego, California, USA) at Beijing Novogene bio Mdt InfoTech Ltd (Beijing, China). To get the high quality clean data, Fastp (Chen et al., 2018) was run to cut and filter the raw reads with default settings. For the 13 *Trapa* species/taxa sequenced, more than 5G clean data for each sample was generated after removing adapters and low quality reads, and the number of clean reads ranged from 5.22 G (*T. mammillifera*) to 6.06 G (*T. bispinosa*). De novo assembly was carried out using the assembler GetOrganelle v1.7 (Jin et al., 2020) with the default setting. The software Geneious primer (Biomatters Ltd., Auckland, New Zealand) was employed to align the contigs and determine the order of the newly assembled plastomes, with *T. bicornis* genome sequence as a reference. All the annotated cp sequence data reported here were deposited in GenBank with accession numbers shown in Table 1.

### 2.3 Annotation and codon usage

We used the genome annotator PGA (Qu et al., 2019) and software GeSeq (Tillich et al., 2017) to annotate protein coding genes, tRNAs and rRNAs, according to the references of *T. quadrispinosa* (MT_941481) and *T. maximowiczii* (NC_037023). Manual correction was carried out to locate the start and stop codons and the boundaries between the exons and introns. With tRNAscan-SE v1.21, BLASTN searches were further performed to confirm the tRNA and rRNA genes (Schattner et al., 2005). The physical maps of cp genomes were generated by OGDRAW (Lohse et al., 2007).

The relative synonymous codon usage (RSCU) was the ratio of the frequency of a particular codon to the expected frequency of that codon, which was obtained by DAMBE v6.04 (Cauzsantos et al., 2017). When the value of RSCU is larger than 1, the codon is used more often than expected. Otherwise, when the RSCU value < 1, the codon is less used than expected (Sharp et al., 1987).

### 2.4 Comparative genomic analyses

Comparative genomic analyses were carried out among the 15 wild *Trapa* species/taxa, which included the 13 species/taxa newly sequenced, and two published ones (*T. kozhevnikovirum* and *T. incisa*) (Wang et al., 2021; Wagutu et al., 2021). The published cp genomes were downloaded from the National Center for Biotechnology Information (NCBI) organelle genome database (https://www.ncbi.nlm.nih.gov).

The mVISTA program in Shuffle-LAGAN mode was used to compare the complete cp genome within the 15 species/taxa, with the annotation of *T. quadrispinos* as a reference (MT_941481). After a manual alignment using MEGA X, all regions, including coding and non-coding regions, were extracted to detect the hyper-variable sites (Kumar et al., 2018). The nucleotide variability (Pi) was computed using DnaSP 5.10 (Librado and Rozas, 2009).

### 2.5 Analysis of repeat sequences and SSRs

Repeat sequences, including forward, palindromic, reverse and complement repeats, were detected by REPuter (Kurtz and Schleiermacher, 1999). The parameters were set with repeat size of ≥30 bp and 90% or greater sequence identity (hamming distance of 3).

Simple sequence repeats (SSRs) were identified using MISA perl script (Thiel et al., 2003), with the threshold number of repeats set as 10, 5, 4, 3, 3, and 3 for mono-, di-, tri-, tetra-, penta-, and hexa-nucleotide SSRs, respectively.

### 2.6 Phylogenetic analyses

Phylogenetic analyses based on the complete chloroplast genomes were carried out for the 18 species/taxa, including the 13 newly sequenced species/taxa and five published ones (*T. bicornis*, *T. natans*, *T. maximowiczii*, *T. insica* and *T. kozhevnikovirum*). Considering the close relationship between Trapaceae and Sonneratiaceae/Lythraceae (Sun et al., 2020), *Sonneratia alba* (Sonneratiaceae) and two Lagerstroemia species (*L. calyculata* and *L. intermedia*, Lythraceae) were included as outgroups. The whole cp genome of the five published *Trapa* species and the three species as outgroups were downloaded from Genbank.

The sequences were aligned using program Mafft 7.0 (Katoh and Standley, 2013) with default parameters. Then, the F81+G was selected as the best-fit model of nucleotide substitution by software Jmodeltest 2 (Darriba et al., 2012). The phylogenetic trees were constructed using three methods: (1) A maximum likelihood (ML) tree was performed using PhyML v.3.0 (Guindon et al., 2010) with 5000 bootstrap replications. The result was visible by the software Figtree v1.4 (https://github.com/rambaut/figtree/releases); (2) The Maximum Parsimony (MP) tree was run by the mega X (Kumar et al., 2018) with 5000 bootstrap values; (3) Bayesian inference (BI) tree was built by the MrBayes v. 3.2.6 (Ronquist et al., 2012) with 2,000,000 generations and sampling every 5000 generations. The first 25% of all trees were regarded as “burn-in” and discarded, and the Bayesian posterior probabilities (PP) were calculated from the remaining trees.

## 3. Results and discussion

### 3.1 Structure and content of Chloroplast Genome

Previous studies showed that the length of cp genome ranged from 120-170kb (Ruhlman and Jansen, 2014). In the present study, the cp genomes of the 15 *Trapa* species/taxa have medium size with 155,453-155,559 bp, compared to the recent study from Lythraceae with 152,049-160,769 bp (Xu et al., 2017; Zheng et al., 2020). The two species with small seeds showed the smallest cp genome size with 155, 453 for *T. maximowiczii* and 155, 477 for *T. incisa*. For the 13 *Trapa* species/taxa with large seeds, the cp genomes of the three species/taxa (*T. bispinosa*, *T. quadrispinosa* and *T. macropoda* var. *bispinosa*) showed the relative small size with 155, 485-155, 495 bp, and the others exhibited similar size with 155, 535 bp (*T. litwinowii*) - 155, 559 bp (*T. mammillifera* and *T. natans* var. *baidangensis*) (Table 2).

Unlike those of the non-photosynthetic (parasitic and mycoheterotrophic) plants, the plastid genomes of green plants are usually highly conserved with the quadripartite structure (Naumann et al., 2016). The typical quadripartite structure was also found in *Trapa*, which consisted of a pair of inverted repeat regions (IRs) (26, 380 – 26, 388 bp) separated by the small single copy (SSC) (18, 265 – 18, 279 bp) and large single copy (LSC) regions (88, 398 – 88, 512 bp). This suggested the difference in cp genome size of wild *Trapa* species mainly occurred in LSC region.

The 15 *Trapa* speices/taxa showed the identical level of GC content with the total content 36.40-36.41%, which was close to that of *Lagerstroemia* (37.57 - 37.72%, Zheng et al., 2020) and Lythraceae (36.41 - 37.72%, Gu et al., 2019). Previous studies detected that the GC content varied greatly among different regions of cp genome. High level of GC content was generally contained in rRNA genes which were located in IR regions (Zheng et al., 2020). Similarly, for *Trapa*, the GC content of rRNA genes reached to 55.48%, which resulted in the highest GC content in the IR regions (42.77%) compared with that of the other two regions (LSC, 34.17 - 34.20%; SSC, 30.17 - 30.21%) (Figure 1; Table 4).

**Figure 1.**
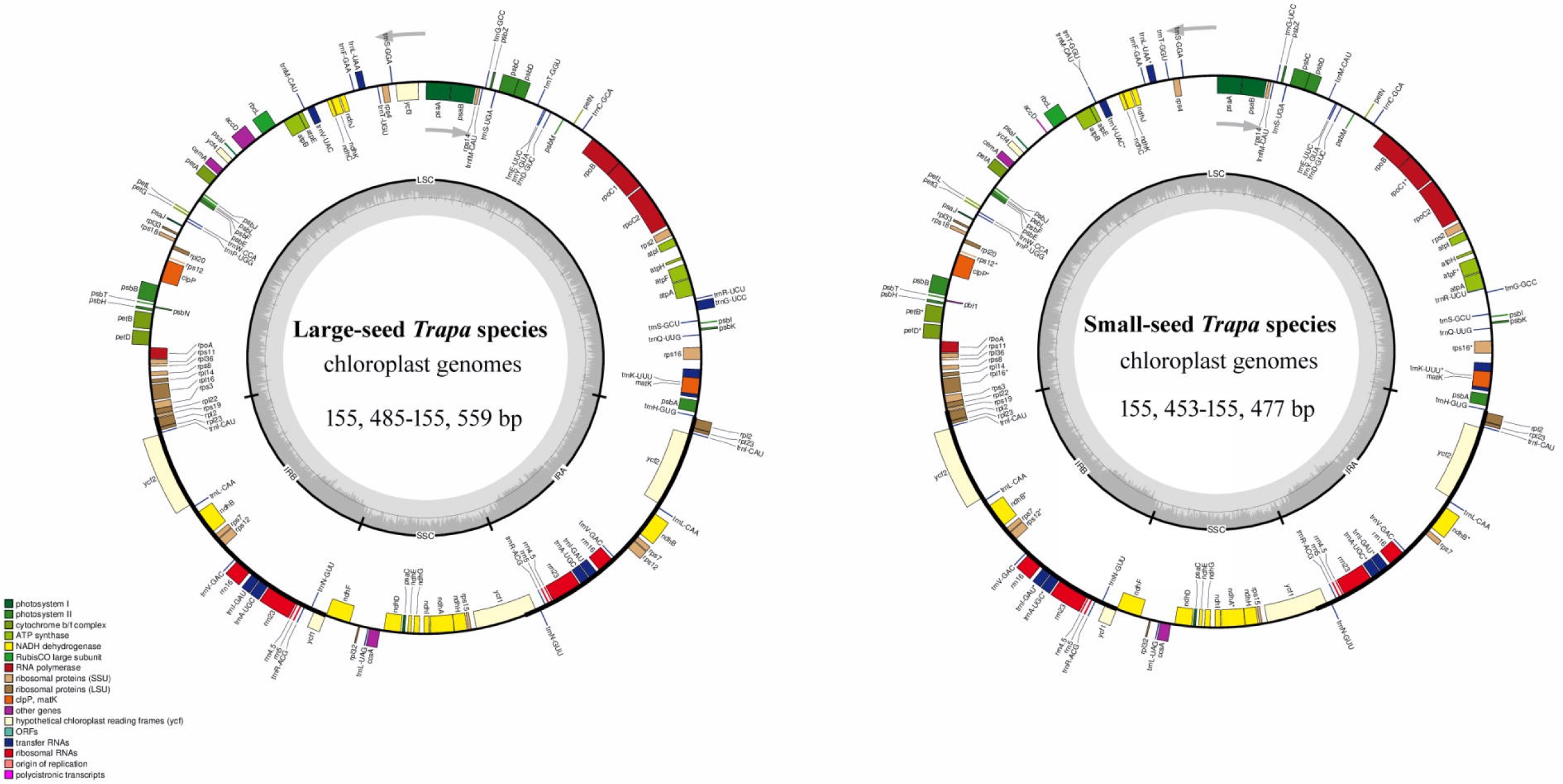
Structural map of the *Trapa* chloroplast genome. Genes drawn inside the circle are transcribed clockwise, and those outside are counterclockwise. Small single copy (SSC), large single copy (LSC), and inverted repeats (IRa, IRb) are indicated. Genes belonging to different functional groups are color-coded.

**Figure 2.**
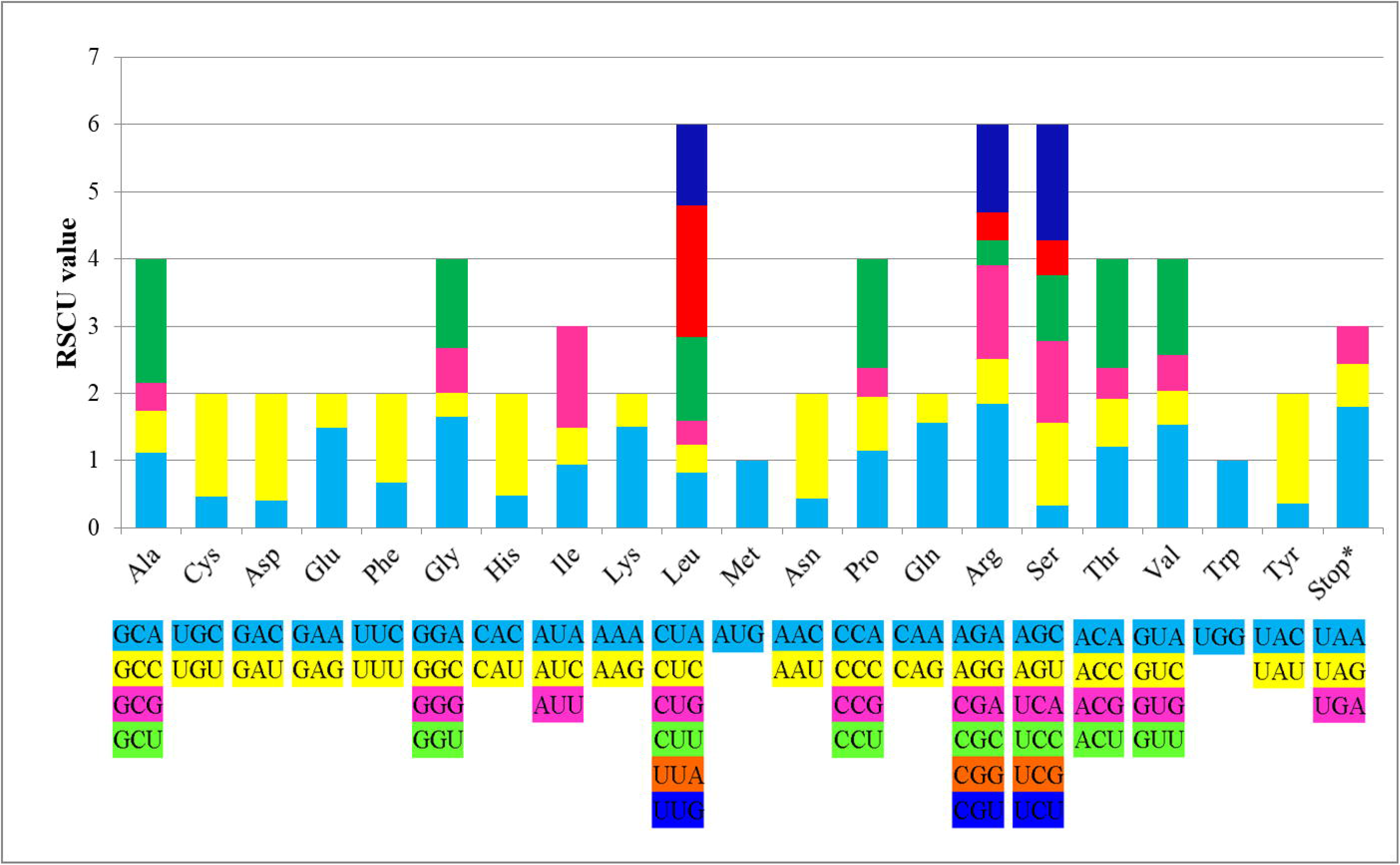
Codon contents of 20 amino acids and stop codon of coding genes of *Trapa* chloroplast genome. Color of the histogram corresponds to the color of codons.

There were 130 genes annotated in the cp genomes of the 13 *Trapa* species/taxa with large seeds, compared with the 129 genes in the two small-seed *Trapa* species (*T incisa* and *T. maximowiczii*). Previous studies showed that the number of PCGs and tRNAs varied in different species, but the number of rRNA was stable (Zhang et al., 2020b), which was proved in the present study. The cp genomes of the 13 *Trapa* species/taxa with large seeds contained 85 protein-coding genes (PCGs), 37 tRNA genes and 8 rRNA genes. In comparison, the two *Trapa* species with small seeds had 83 PCGs, 38 tRNA genes and 8 rRNA genes. The two small-seed species lost the gene *ycf3* related to the conserved open reading frames and one *rps12* for self-replication. Among the unique genes, 44 genes were related to photosynthesis and 59 genes were associated with self-replication (Table 3).

### 3.2 Hypervariable Regions and Comparative Genomic Analysis

Except the gene space, the remaining regions were non-coding sequences, comprising intergenic spacers (IGS) regions, introns and pseudogenes (Lu et al., 2017). In this study, the intergenic regions and introns among the 15 *Trapa* species/taxa ranged from 46, 592 to 46, 812 bp and from 15, 957 to 18, 852 bp, respectively. IGS and introns generally have higher nucleotide substitution rates than the protein-coding regions, which yield many phylogenetically informative sites (Shaw et al. 2007; Jansen and Ruhlman, 2012). In this study, high level of the nucleotide diversity (Pi) values was found in the non-coding regions (0 - 0.0123, with average 0.000857) compared with those of the coding regions (0 - 0.00282, with average 0.000217) (Figure 7).

Previous studies exhibited a negative correlation between the GC content and the variability of cp genome sequences (Zhang et al., 2020a; Zheng et al., 2020). This was proved in the present study. Most variable regions for *Trapa* were located in IGS regions with the lowest GC content (30.28-30.42%) (Table 4). And the highest Pi values (0.00416-0.0123) were found in the IGS regions, including *ndhE—ndhG*, *psbA—trnK-UUU*, *psbK—psbI*, *rps2—rpoC2* and *atpA—atpF* (Figure 7). Therefore, IGS is often used as DNA barcoding markers in many species (Nguyen et al., 2015). It is worth noting that IGS regions were mostly distributed in the LSC region. Among the 20 highest Pi values, 16 divergence hotspots with low sequence identity were found, which included 11 regions (*psaC*, *clpP*, *psbE*, *atpF*, *rps2*—*rpoC2*, *psbK*—*psbI*, *psbA*—*trnK-UUU*, *trnS-GCU*—*trnG-UCC*, *trnH-GUG*—*psbA*, *trnT-GGU*—*psbD* and *accD-psaI*) in the LSC and 5 regions (*ndhD*, *ndhI*, *ycf1*, *ndhF* and *ndhE*—*ndhG*) in the SSC (Figure 3).

**Figure 3.**
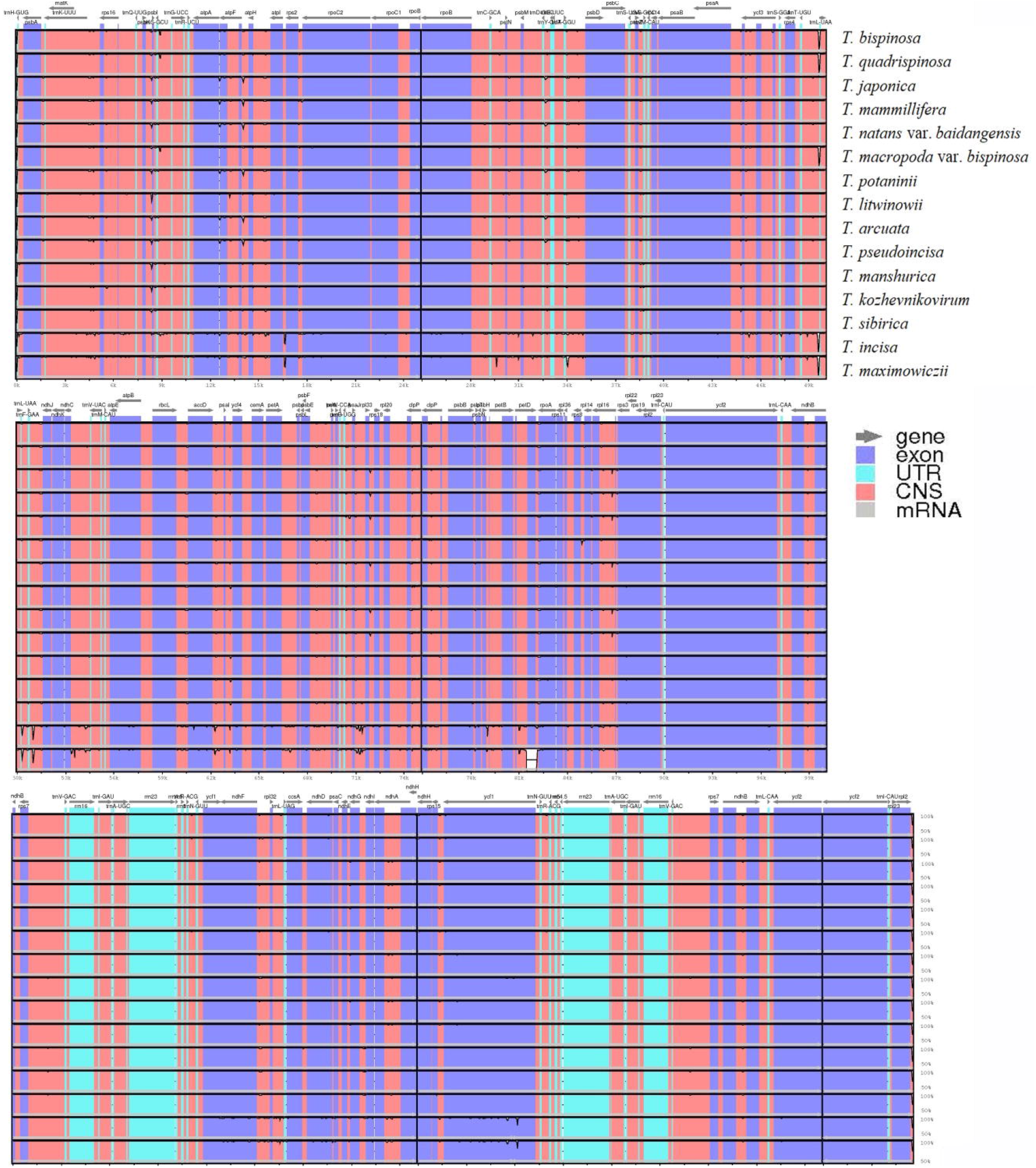
Sequence alignment of whole chloroplast genomes using the Shuffle LAGAN alignment algorithm in mVISTA. *Trapa bicornis* was chosen to be the reference genome. The vertical scale indicates the percentage of identity, ranging from 50 to 100%.

**Figure 4.**
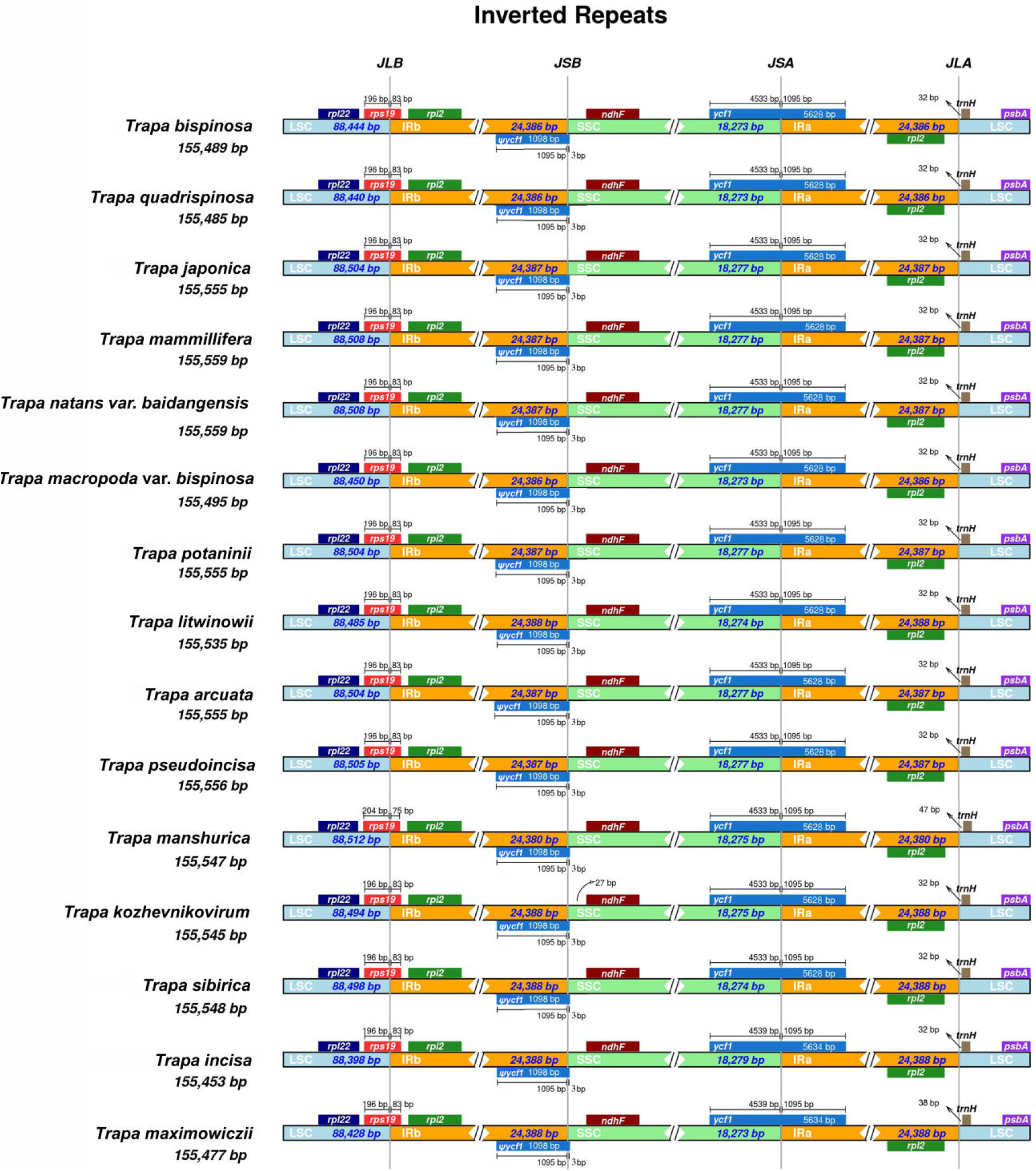
Comparison of junctions between the LSC, SSC, and IR regions among 15 species. Distance in the figure is not to scale. LSC Large single-copy, SSC Small single-copy, IR inverted repeat.

We found 15/16 genes containing one or two introns for the small-seed/large-seed species, consisting of 6 tRNA genes (*trnK-UUU*, *trnA-UGC*, *trnI-GAU*, *trnG-UCC*, *trnV-UAC* and *trnL-UAA*) and 8 protein-coding genes (PCGs) (*rps16*, *rpoC1*, *atpF*, *ndhB*, *ndhA*, *petD*, *petB and rpl16*) with one intron, and one PCG *clpP* with two introns for the small-seed *Trapa* species, but two PCGs (*ycf3* and *clpP*) for the large-seed ones. The introns of the PCGs shared the similar position and length in each of the 15 *Trapa* taxa (Table 5), suggesting the high conservation of cp genome within *Trapa*. The *rpl2* intron loss was considered as one of the important evolutionary event in Lythraceae, which was presumed to occur after the divergence of *Lythraceae* from Onagraceae (Chao et al., 2017; Gu et al., 2016, 2019; Zheng et al., 2020). Like Lythraceae, the 15 *Trapa* species/taxa exhibited the loss of *rpl2* intron, which might suggest a close genetic relationship between Lythraceae and Trapaceae.

The gene contains several internal stop codons, which tends to be a pseudogene (Lu et al., 2017; Sloan et al., 2014). Alternatively, if the sequence was conserved over broad evolutionary distances and lack of internal stop, it tends to be assumed as functional protein-coding genes (Raubeson et al., 2007). In the present study, though the *ycf3* gene was only detected in the large-seed *Trapa* taxa, none internal stop codon was found within the gene. Therefore, instead of a pseudogene, the *ycf3* appears as a protein-coding gene.

### 3.3 Repeat Structure and cpSSR

For cp genomes, long repeats play an important role in the sequence rearrangement and recombination, which could be as modules of large assemblies and participate in catalytic function or protein binding (Nazareno et al. 2015; Zheng et al., 2020). A total of 595 long repetitive repeats (30 - 65bp) were identified from the 15 *Trapa* taxa, consisting of 324 forward, 230 palindromic, 25 reverse and 16 complementary repeats. For the genus *Trapa*, the size of top three most frequently shown long repeats was 30bp, 31bp and 65bp, which was found 227, 77 and 60 times, respectively. For each species/taxon, the number of long repeats varied from 37 (*T. incisa* and *T. maximowiczii*) to 41 (*T. sibirica*); and the number of forward, palindromic, reversed and complementary repeats were 19-24, 15-16, 0-3 and1-2, respectively (Figure 5). Additionally, most long repetitive repeats were distributed in intergenic areas, and a few existed in shared genes, such as *ycf2*, which was analogous to the results of other angiosperm lineages (Yang et al., 2017; Zheng et al., 2020).

**Figure 5.**
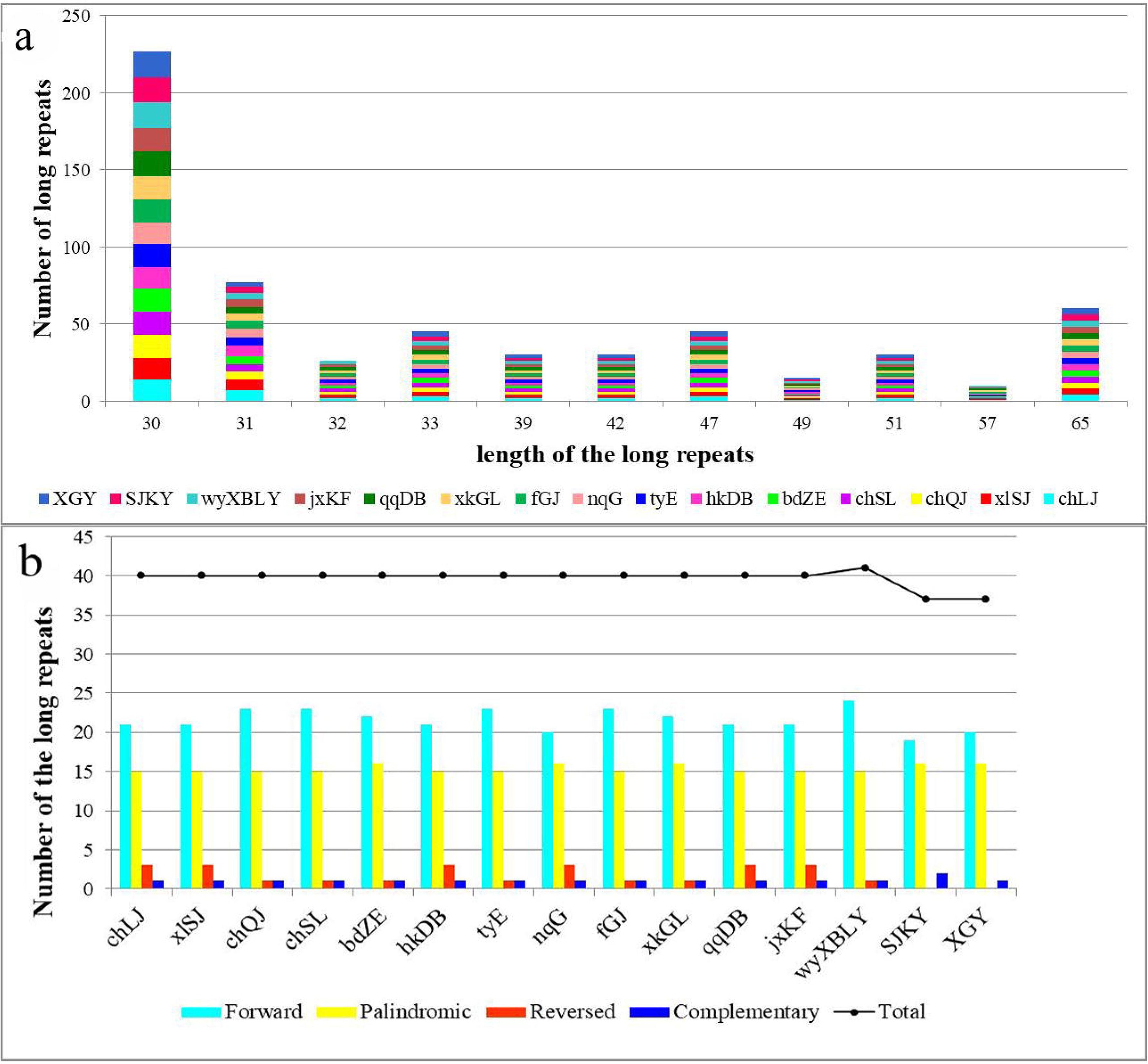
Number of long repetitive repeats on the complete chloroplast genome sequence of 15 *Trapa* species. (a) frequency of the repeats more than 30 bp, (b) frequency of repeat types.

Due to a high polymorphism rate at the intra- and inter-species level, cp genome SSRs have been viewed as excellent molecular markers in population genetics and phylogenetic research (Xue et al., 2017; Zheng et al., 2020). In this study, for each species/taxon, the number of total SSRs was from 110 (*T. quadrispinosa*) to 123 (*T. incisa* and *T. maximowiczii*). Most cp SSRs, from 83.48% (*T. japonica*, *T. manshurica* and *T. litwinowii*) to 86.18% (*T. maximowiczii*), were distributed in the LSC regions, which was higher than that in the genus *Lagerstroemia* (66.55%; Xu et al., 2017) and *Myrsinaceae* (74.37%; Yan et al., 2019). Among these SSRs, the mononucleotide A/T repeat units occupied the highest proportion with 78.18 - 80.49%, and the second and third highest proportions were dinucleotide repeats (AC/GT and AT/AT) and tetranucleotide repeats (AAAG/CTTT, AAAT/ATTT, AACC/GGTT, AAGT/ACTT, AATT/TTAA, ACAT/ATGT and AGAT/ATCT) accounting for 9.40 - 10.00% and 8.13 - 9.09%, respectively (Figure 6). The observed high AT content in cp SSRs was also found in the other genera (Asaf et al., 2016; Chao et al., 2017; Kuang et al., 2011)

**Figure 6.**
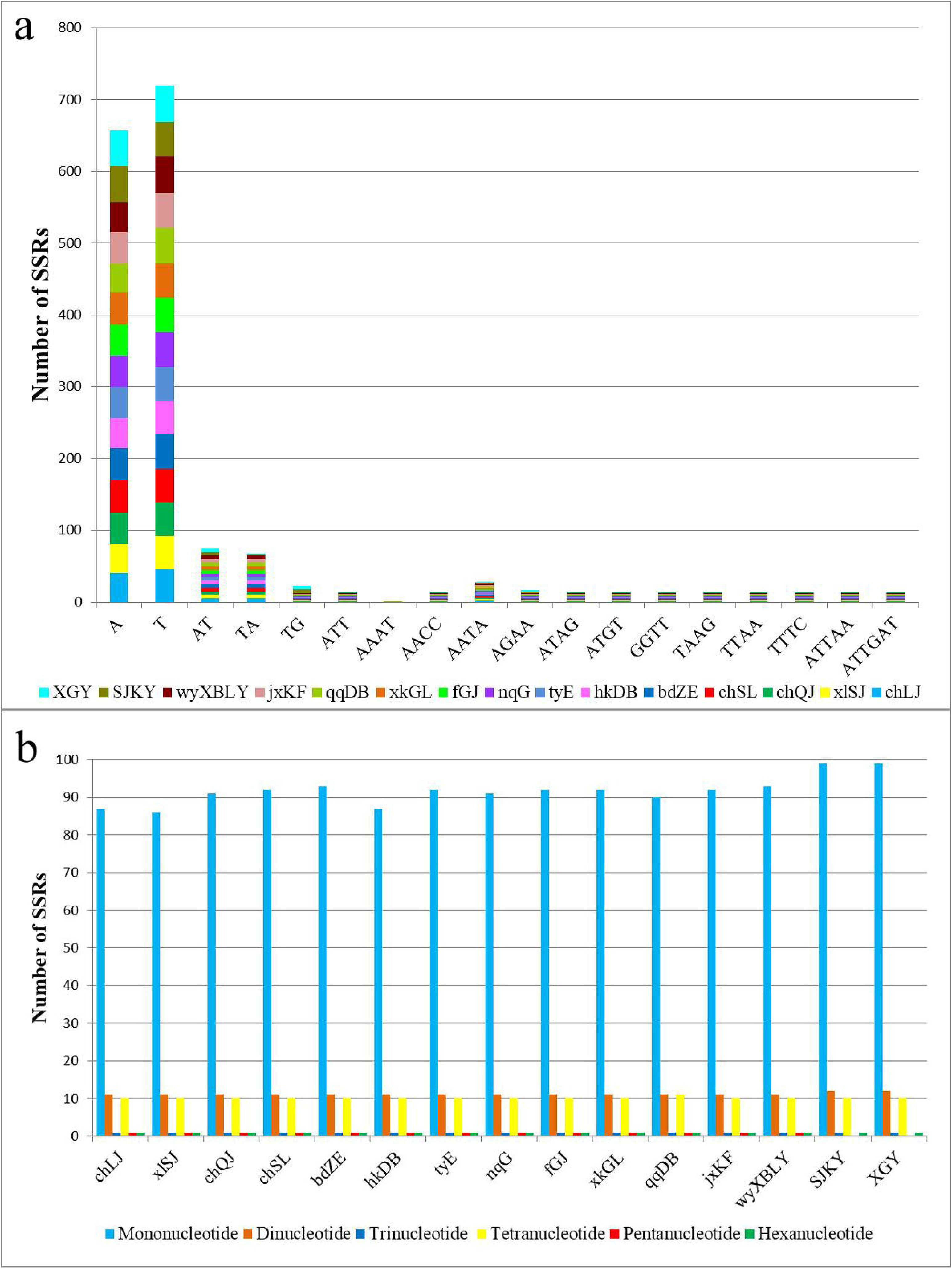
The comparison of simple sequence repeats (SSRs) distribution in 15 chloroplast genomes. (a) frequency of common motifs; (b) number of different SSR types.

**Figure 7.**
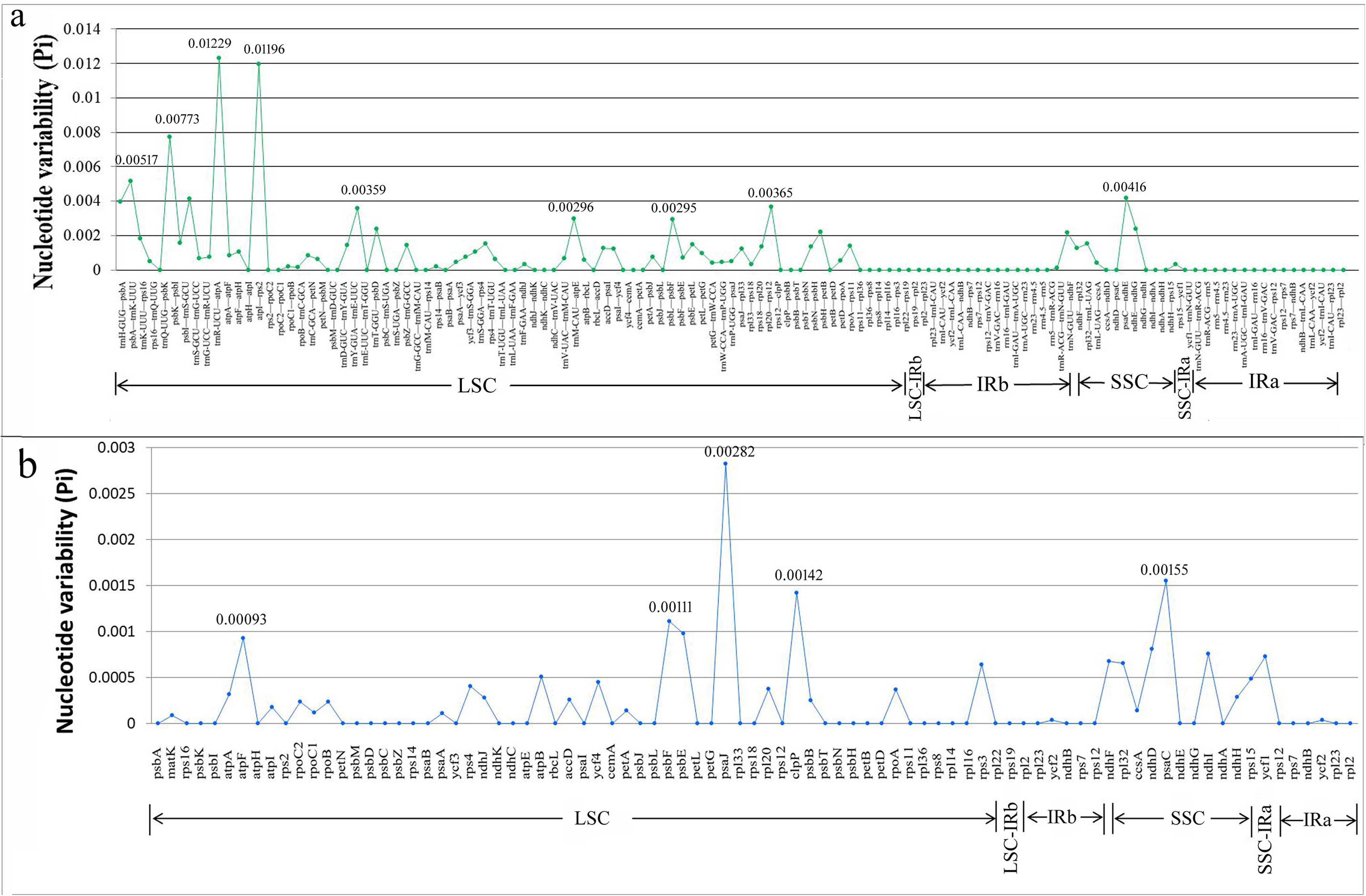
The nucleotide variability (Pi) value in the 15 *Trapa* chloroplast genomes. (a) Intergenic regions. b Protein-coding genes. These regions are arranged according to their location in the chloroplast genome.

### 3.4 Boundaries of IR regions and codon usage

For angiosperms, the boundaries between IR regions and single-copy (SC) regions result in the difference of genome size by expansion or shrinkage (Kim and Lee, 2004; Lin et al., 2012). In this study, the junctions of IR/SC regions were compared among the 15 *Trapa* species\taxa. The cp genomes of the 15 *Trapa* plants showed many minor differences in the boundary regions although the number and order of genes were highly conserved. The *rps*19 gene spanned the LSC/IRb border and extended into the IRb region with the length 75-83 bp (except *T. manshurica* with 75 bp, the others were 83 bp). The *ycf*1 gene covered the junction of SSC/IRa, and the part of *ycf*1 into IRa region showed a stable size (1095 bp) for all the *Trapa* plants. On the contrary, the remaining part of *ycf*1, located in SSC, showed a variable size, with 4539 bp for the two small-seed species and 4533 bp for the remaining 13 taxa. Accordingly, a pseudogene fragment ψ*ycf*1 was detected in the border of IRa/LSC with 1098 bp for all *Trapa* taxa, and the parts located in IRb and SSC were 1095 bp and 3 bp, respectively. The gene *trnH* was distributed on the right side of the border of IRb/LSC, with an interval of 32 - 47 bp from the border to the gene (Figure 4).

For all the *Trapa* species/taxa, a total of 64 types of codons encoding 20 amino acids were detected. In all, the 83/85 protein coding genes within *Trapa* encoded 25, 832 to 26, 590 codons. The results were comparable to that of the genus *Lagerstroemia*, 79 genes encoding 25, 068 - 27, 111 codons in their cp genomes (Zheng et al., 2020). Three termination codons, UGA, UAG and UAA, were found for all the *Trapa* species. The highest number of codon occurrences (> 1000) was found in GAA, UUU, AUU and AAA, which encoded Glutamicacid, Phenylalanine, Isoleucine and Lysine, respectively. Leucine, Isoleucine and Serine were the top three amino acids with high frequency, , with the frequency 2771-2826, 2229-2298 and 1992-2067, respectively. Conversely, Cysteine (288-298) and Tryptophan (459-468) were the least frequent amino acids.

Like that of *Lagerstroemia* (Zheng et al., 2020), the RSCU value of a single amino acid showed a positive correlation with the number of codons encoding it. The highest and lowest RSCU values were exhibited in the UUA encoding Leucine (1.96) and the AGC encoding Serine (approximately 0.34), respectively (Figure 2). The results also showed that 30 codons were used frequently with RSCU values > 1, and all of them ended with A/U, which might be associated with the high proportion of A/T in cp genomes (Eguiluz et al., 2017).

### 3.5 Phylogenetic Analysis

Based on the whole cp genomes, the three phylogenetic trees (MP, ML and BI) showed the similar clustering patterns. The species from the same family clustered together. At first, the branch with the two *Lagerstroemia* species (Lythraceae) separated from the other species. And *S. alba* (Sonneratiaceae) showed to be a sister of *Trapa* species, which was also supported by recent studies (Sun et al., 2020; Wang et al., 2021; Wagutu et al., 2021).

All the *Trapa* species/taxa were divided into two clusters, the clusters of small-seed species and large-seed species, which were consistent with the results of morphological studies (Fan et al., 2016). The two species with small seeds initially separated from the other *Trapa* taxa, which agreed to the results of AFLPs study recently published (Fan et al., 2021). Additionally, the present study further detected a close genetic relationship between the two species with small seeds. On the other hand, in the cluster with large seeds, the cultivated species *T. bicornis* was the outermost part of the group, which might suggest a complex origin of cultivated *Trapa* species. However, it was reported that another common cultivated species *T. bicornis* var. *cochinchinensis* demonstrated a hybrid origin between *T. quadrispinosa* and *T. bispinosa* (Fan et al., 2021). The remaining large-seed *Trapa* taxa were divided into three clusters: (1) the first cluster included three *Trapa* species (*T. quadrispinosa*, *T. bispinosa* and *T. macropoda* var. *bispinosa*). The intimate relationship between the former two has been proved by many studies (Li et al., 2017; Fan et al., 2021). It was the first time for *T. macropoda* var. *bispinosa* being reported, and the only difference between this species and *T. bispinosa* was the larger seed bottom of this species; (2) the second cluster contained six species/taxa (*T. japonica*, *T. mammillifera*, *T. potaninii*, *T. pseudoincisa*, *T. arcuata* and *T. natans var. baidangensis*), which were of the deformed and obvious tubercles and thick husks; (3) the third cluster covered four species (*T. litwinowii*, *T. manshurica*, *T. kozhevnikovirum* and *T. sibirica*), which possess smooth and tight seed coats and were collected from the Amur River (Figure 8). Among them, *T. litwinowii* is the only one with two horns. Compared with *T. litwinowii*, *T. manshurica* shares the same seed size and seed ornamentations, but *T. manshurica* has four horns. *T. manshurica* and *T. sibirica* share similar morphologic characteristics, but *T. sibirica* has larger nuts, small crown and longer fruit neck. The seed crown of the three *Trapa* species (*T. litwinowii*, *T. manshurica* and *T. sibirica*) was large and curled outwardly. *T. kozhevnikovirum* and *T. sibirica* have the similar seed size, but *T. kozhevnikovirum* has a small seed crown and unobvious seed neck.

**Figure 8.**
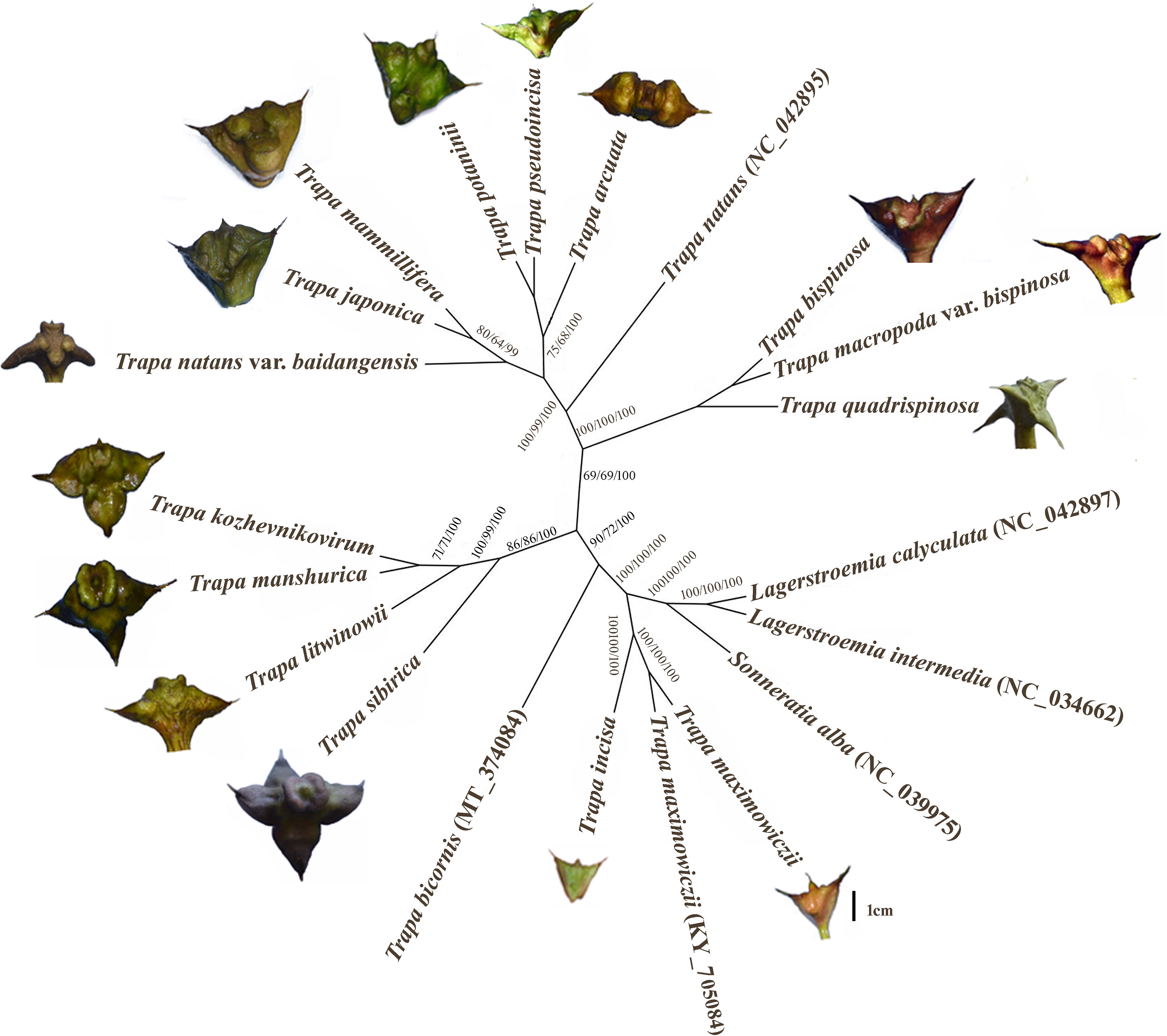
The phylogenetic tree is based on 21 complete chloroplast genome sequences using Maximum likelihood (ML), Maximum parsimony (MP) and Bayesian inference (BI). The number above the lines indicates bootstrap values for ML and MP and posterior probabilities for BI of the phylogenetic analysis for each clade.

## 4. Conclusions

This study reported the comparative analysis results of 15 *Trapa* cp genome sequences with detailed gene annotation. The 15 cp genomes are of the similar quadripartite structure with a high degree of the synteny in gene order, suggesting the high conservation of the cp genome in *Trapa*. There was no obvious change found in the IR/SC junctions, indicating the junctions were relatively conservative in *Trapa*. Most of the long repetitive repeats and SSRs were detected in the intergenic and LSC regions, which provided effective cp markers for species identification, genetic relationship analysis and the population genetics. Three phylogenetic analysis approaches (MP, ML and BI methods) consistently supported the 15 *Trapa* taxa separated into two major evolutionary branches, large- and small-seed branches, which was agreeable to the results of morphological studies. And a close genetic relationship was found between the two small-seed species. The baseline cp genome information of *Trapa* should be useful for species identification and phylogeographic analysis of *Trapa* species/taxa, which will be beneficial to the management and utilization of genetic resources of the genus.

## Supporting information

Tables for the manuscript

## Author contributions

FX collected and identified the species of sample, designed the experiments, analyzed the data and wrote the paper. WW performed the experiments, contributed reagents/materials/analysis tools, prepared figures and/or tables and wrote the paper. CY conceived and designed the experiments, reviewed drafts of the paper. GKW contributed reagents/materials/analysis tools. LW and LX help conceived and designed the experiments and reviewed the paper.

