## Supplementary material for "Comparative analyses and phylogenetic relationships of 15 *Trapa* (Trapaceae) species based on Complete Chloroplast Genomes": Tables for the manuscript: cptables0331.docx

| **No.** | **Abbr.** | **Species** | **Location** | **Voucher No.** | **GenBank No.** |
| --- | --- | --- | --- | --- | --- |
| 1 | chLJ | *Trapa bispinosa* Roxb. | Changhu Lake, Hubei | yychen20180060 | MW579848 |
| 2 | xlSJ | *Trapa quadrispinosa* Roxb. | Xiliang Lake, Hubei | yychen20180055 | MW037838 |
| 3 | chQJ | *Trapa japonica* Flerow | Changhu Lake, Hubei | yychen20180061 | MW579849 |
| 4 | chSL | *Trapa mammillifera Miki* | Changhu Lake, Hubei | yychen20180059 | MW579850 |
| 5 | bdZE | *Trapa natans* var. *baidangensis* | Baidang Lake, Anhui | yychen20180097 | MW784170 |
| 6 | hkDB | *Trapa macropoda* Miki *var. bispinosa* W. H. Wan | Haikou Lake, Hubei | yychen20180075 | MW579851 |
| 7 | tyE | *Trapa potaninii* V. Vassil | Tangyuan, Heilongjiang | yychen20180019 | MW579852 |
| 8 | nqG | *Trapa litwinowii* V. Vassil | Nongqiao, Heilongjiang | yychen20180034 | MW579853 |
| 9 | fGJ | *Trapa arcuata* S. H. Li et Y. L. Chang | 856farm, Heilongjiang | yychen20180046 | MW579854 |
| 10 | xkGL | *Trapa pseudoincisa* Nakai | Xunke, Heilongjiang | yychen20180007 | MW579855 |
| 11 | qqDB | *Trapa manshurica* Fler. | Qiqihaer, Heilongjiang | yychen20180001 | MW579857 |
| 12 | jxKF | *Trapa kozhevnikovirum* Pshennikova | Jixi, Heilongjiang | yychen20180042 | MW027640 |
| 13 | wyXBLY | *Trapa sibirica* Fler. | Wuyun, Heilongjiang | yychen20180018 | MW579856 |
| 14 | SJKY | *Trapa incisa* Sieb. et Zucc. | Wuhan Botanical Garden, Hubei | yychen20180066 | MW543307 |
| 15 | XGY | *Trapa maximowiczii* Korsch． | Tangyuan, Heilongjiang | yychen20180023 | MW579858 |

Table 1 The GenBank accession numbers of 15 species using in phylogenetic analysis

Table 2 Summary of complete chloroplast genomes for 15 *Trapa* species

| **Taxon** | **chLJ** | **xlSJ** | **chQJ** | **chSL** | **bdZE** | **hkDB** | **tyE** | **nqG** | **fGJ** | **xkGL** | **qqDB** | **jxKF** | **wyXBLY** | **SJKY** | **XGY** |
| --- | --- | --- | --- | --- | --- | --- | --- | --- | --- | --- | --- | --- | --- | --- | --- |
| **Total length(bp)** | 155,489 | 155,485 | 155,555 | 155,559 | 155,559 | 155,495 | 155,555 | 155,535 | 155,555 | 155,556 | 155,547 | 155,545 | 155,548 | 155,477 | 155,453 |
| **GC(%)** | 36.41 | 36.41 | 36.40 | 36.40 | 36.40 | 36.41 | 36.40 | 36.40 | 36.40 | 36.40 | 36.41 | 36.41 | 36.40 | 36.41 | 36.41 |
| **Length of LSC(bp)** | 88,444 | 88,440 | 88,504 | 88,508 | 88,508 | 88,450 | 88,504 | 88,485 | 88,504 | 88,505 | 88,512 | 88,494 | 88,498 | 88,428 | 88,398 |
| **GC of LSC(%)** | 34.18 | 34.18 | 34.18 | 34.18 | 34.18 | 34.18 | 34.17 | 34.18 | 34.18 | 34.18 | 34.18 | 34.18 | 34.17 | 34.19 | 34.20 |
| **Length of SSC(bp)** | 18,273 | 18,273 | 18,277 | 18,277 | 18,277 | 18,273 | 18,277 | 18,274 | 18,277 | 18,277 | 18,275 | 18,265 | 18,274 | 18,273 | 18,279 |
| **GC of SSC(%)** | 30.20 | 30.21 | 30.18 | 30.18 | 30.18 | 30.20 | 30.17 | 30.19 | 30.17 | 30.17 | 30.19 | 30.20 | 30.18 | 30.17 | 30.17 |
| **Length of IR(bp)** | 24,386 | 24,386 | 24,387 | 24,387 | 24,387 | 24,386 | 24,387 | 24,388 | 24,387 | 24,387 | 24,380 | 24,388 | 24,388 | 24,388 | 24,388 |
| **GC of IR(%)** | 42.77 | 42.77 | 42.77 | 42.77 | 42.77 | 42.77 | 42.77 | 42.77 | 42.77 | 42.77 | 42.77 | 42.77 | 42.77 | 42.77 | 42.77 |
| **Total number of genes** | 130 | 130 | 130 | 130 | 130 | 130 | 130 | 130 | 130 | 130 | 130 | 130 | 130 | 129 | 129 |
| **Protein-coding gene** | 85 | 85 | 85 | 85 | 85 | 85 | 85 | 85 | 85 | 85 | 85 | 85 | 85 | 83 | 83 |
| **tRNA** | 37 | 37 | 37 | 37 | 37 | 37 | 37 | 37 | 37 | 37 | 37 | 37 | 37 | 38 | 38 |
| **rRNA** | 8 | 8 | 8 | 8 | 8 | 8 | 8 | 8 | 8 | 8 | 8 | 8 | 8 | 8 | 8 |

Table 3 Genes in the sequenced *Trapa* chloroplast genome

| **Category of genes** | **Function of genes** | **Name of genes** |
| --- | --- | --- |
| Subunits of ATP synthase | Genes for photosynthesis | *atp A, B, E, F, H, I* |
| Other genes |  | *accD, ccsA,* *cemA, clpP, matK* |
| Subunits of NADH dehydrogenase | Genes for photosynthesis | *ndh A, B(2), C, D, E, F, G, H, I, K, J* |
| Subunits of photosystem I |  | *Psa A, B, C, I, J* |
| Subunits of photosystem II | Genes for photosynthesis | *psb A, B, C, E, F, G, H, I, J, K, L, M, N, T, Z* |
| Subunits of cytochrome |  | *pet A, B, D, G, L, N* |
| Large subunit of Rubisco | Genes for photosynthesis | *rbcL* |
| Large subunit of ribosome | Self-replication | *rps 2(2), 3, 4, 7(2), 8, 11, 12(2)^A^, 14, 15, 16, 18, 19* |
| DNA dependent RNA polymerase | Self-replication | *rpo A, B, C1, C2* |
| Ribosomal RNA genes | Self-replication | *rrn* 5(2)*,*4.5(2), 16(2), 23(2) |
| Small subunit of ribosome | Self-replication | *rpl*2(2), 14, 16, 20, 22, 23(2), 32, 33, 36 |
| Transfer RNA genes | Self-replication | *trnA-UGC*(2)*, trnC-GCA, trnD-GUC, trnE-UUC, trnF-GAA, trnfM-CAU, trnG-GCC, trnG-UCC, trnH-GUG, trnI-CAU*(2), *trnI-GAU*(2), *trnK-UUU, trnL-CAA*(2)*, trnL-UAA, trnL-UAG, trnM-CAU*(x2)*, trnN-GUU*(2)*, trnP-UGG, trnQ-UUG, trnR-ACG*(2)*, trnR-UCU, trnS-GCU, trnS-GGA, trnS-UGA, trnT-GGU, trnT-UGU, trnV-GAC*(2), *trnV-UAC, trnW-CCA, trnY-GUA* |
| Conserved open reading frames | Genes of unknown function | *ycf 1(2), 2(2), 3^A^, 4* |

*^A^* indicates the loss of the gene in the two small-seed *Trapa* species; (2) indicates the gene duplicated in all species; (x2) indicates the gene just duplicated in the two small-seed *Trapa* species.

Table 4 Distribution of genes and intergenic regions for 15 *Trapa* species

| Species | Protein Coding Genes | |  | rRNA | |  | tRNA | |  | Intergenic Regions | |  | Intron | |
| --- | --- | --- | --- | --- | --- | --- | --- | --- | --- | --- | --- | --- | --- | --- |
|  | Length(bp) | GC(%) |  | Length(bp) | GC(%) |  | Length(bp) | GC(%) |  | Length(bp) | GC(%) |  | Length(bp) | GC(%) |
| chLJ | 78672 | 37.28 |  | 9034 | 55.48 |  | 2812 | 53.31 |  | 46713 | 30.31 |  | 18838 | 36.14 |
| xlSJ | 78672 | 37.28 |  | 9034 | 55.48 |  | 2812 | 53.31 |  | 46713 | 30.31 |  | 18834 | 36.15 |
| chQJ | 78672 | 37.27 |  | 9034 | 55.48 |  | 2812 | 53.31 |  | 46791 | 30.28 |  | 18826 | 36.16 |
| chSL | 78672 | 37.27 |  | 9034 | 55.48 |  | 2812 | 53.31 |  | 46795 | 30.28 |  | 18826 | 36.16 |
| bdZE | 78672 | 37.27 |  | 9034 | 55.48 |  | 2812 | 53.31 |  | 46795 | 30.29 |  | 18826 | 36.16 |
| hkDB | 78672 | 37.28 |  | 9034 | 55.48 |  | 2812 | 53.31 |  | 46723 | 30.30 |  | 18834 | 36.15 |
| tyE | 78672 | 37.27 |  | 9034 | 55.48 |  | 2812 | 53.31 |  | 46791 | 30.28 |  | 18826 | 36.16 |
| nqG | 78672 | 37.27 |  | 9034 | 55.48 |  | 2812 | 53.31 |  | 46770 | 30.31 |  | 18827 | 36.14 |
| fGJ | 78672 | 37.27 |  | 9034 | 55.48 |  | 2812 | 53.31 |  | 46791 | 30.28 |  | 18826 | 36.16 |
| xkGL | 78672 | 37.27 |  | 9034 | 55.48 |  | 2812 | 53.31 |  | 46792 | 30.28 |  | 18826 | 36.16 |
| qqDB | 78672 | 37.27 |  | 9034 | 55.48 |  | 2812 | 53.31 |  | 46774 | 30.30 |  | 18835 | 36.14 |
| jxKF | 78627 | 37.24 |  | 9030 | 55.48 |  | 2788 | 55.44 |  | 46812 | 30.42 |  | 18852 | 36.14 |
| wyXBLY | 78498 | 37.27 |  | 9034 | 55.48 |  | 2812 | 53.31 |  | 46770 | 30.29 |  | 18840 | 36.14 |
| SJKY | 77667 | 37.26 |  | 9034 | 55.48 |  | 2861 | 52.50 |  | 46592 | 30.32 |  | 15957 | 36.15 |
| XGY | 77496 | 37.26 |  | 9034 | 55.48 |  | 2861 | 52.50 |  | 46686 | 30.33 |  | 16031 | 36.15 |

Table 5 Length of introns and complete gene of intron-contained protein-coding genes

| *rps16* | *rpoC1* | *atpF* | *ndhB* | *ndhA* | *petD* | *petB* | *rpl16* | *ycf3* | *clpP* |
| --- | --- | --- | --- | --- | --- | --- | --- | --- | --- |
| 866/1133 | 740/2780 | 766/1321 | 684/2217 | 1059/2151 | 742/1225 | 790/1438 | 1012/1420 | 756/770/2033 | 608/818/2017 |
| 866/1133 | 740/2780 | 766/1321 | 684/2217 | 1059/2151 | 742/1225 | 790/1438 | 1012/1420 | 756/766/2029 | 608/818/2017 |
| 864/1131 | 740/2780 | 767/1322 | 684/2217 | 1060/2152 | 742/1225 | 790/1438 | 1003/1411 | 756/766/2029 | 608/819/2018 |
| 864/1131 | 740/2780 | 767/1322 | 684/2217 | 1060/2152 | 742/1225 | 790/1438 | 1003/1411 | 756/766/2029 | 608/819/2018 |
| 864/1131 | 740/2780 | 767/1322 | 684/2217 | 1060/2152 | 742/1225 | 790/1438 | 1003/1411 | 756/766/2029 | 608/819/2018 |
| 866/1133 | 740/2780 | 766/1321 | 684/2217 | 1059/2151 | 742/1225 | 790/1438 | 1012/1420 | 756/766/2029 | 608/818/2017 |
| 864/1131 | 740/2780 | 767/1322 | 684/2217 | 1060/2152 | 742/1225 | 790/1438 | 1003/1411 | 756/766/2029 | 608/819/2018 |
| 865/1132 | 740/2780 | 758/1313 | 684/2217 | 1058/2150 | 742/1225 | 790/1438 | 1012/1420 | 757/766/2030 | 608/820/2019 |
| 864/1131 | 740/2780 | 767/1322 | 684/2217 | 1060/2152 | 742/1225 | 790/1438 | 1003/1411 | 756/766/2029 | 608/819/2018 |
| 864/1131 | 740/2780 | 767/1322 | 684/2217 | 1060/2152 | 742/1225 | 790/1438 | 1003/1411 | 756/766/2029 | 608/819/2018 |
| 865/1132 | 740/2780 | 766/1321 | 684/2217 | 1059/2151 | 742/1225 | 790/1438 | 1011/1419 | 757/766/2030 | 608/820/2019 |
| 869/1136 | 740/2801 | 767/1322 | 684/2163 | 1059/2151 | 742/1225 | 790/1438 | 1012/1420 | 757/766/2030 | 608/820/2019 |
| 865/1132 | 740/2780 | 768/1323 | 684/2217 | 1058/2150 | 742/1225 | 790/1438 | 1013/1421 | 757/768/2032 | 608/820/2019 |
| 865/1132 | 740/2780 | 768/1323 | 685/2218 | 1059/2151 | 751/1234 | 790/1438 | 1011/1419 | - | 605/819/2015 |
| 865/1132 | 740/2780 | 768/1323 | 685/2218 | 1057/2149 | 751/1234 | 790/1438 | 1014/1422 | - | 606/820/2017 |
